# Exploration of clinical breakpoint of Danofloxacin for *Glaesserella parasuis* in plasma and in PELF

**DOI:** 10.1101/2021.04.15.440097

**Authors:** Zihui Xu, Anxiong Huang, Xun Luo, Peng Zhang, Lingli Huang, Xu Wang, Kun Mi, Shiwei Fang, Xiao Huang, Jun Li, Zonghui Yuan, Haihong Hao

## Abstract

**Background:** To establish the clinical breakpoint (CBP) of danofloxacin to *G. parasuis*, three cutoff values, including epidemiological cutoff value (ECV), pharmacodynamic cutoff value (CO_PD_) and clinical cutoff value (CO_CL_), was obtained in the present study.

**Methods:** The ECV was calculated using ECOFFinder base on MIC distribution of 347 *G. parasuis* collected from disease pigs. The CO_PD_ was established base on *in vivo* and *ex vivo* pharmacokinetic (PK) - pharmacodynamic (PD) modeling of danofloxacin both in plasma and pulmonary epithelial lining fluid (PELF) using Hill formula and Monte Carlo analysis. The CO_CL_ was established based on the relationship between possibility of cure (POC) and MIC in the clinical trials using “WindoW” approach, nonlinear regression and CART analysis.

**Results:** The MIC_50_ and MIC_90_ of danofloxacin against 347 *G. parasuis* were 2 μg/mL and 8 μg/mL, respectively. The ECV value was set up as 8 μg/mL using ECOFFinder. Concentration-time curve of danofloxacin indicated a two-compartment model for PK analysis. The PK parameters of the maximum concentration (C_max_) and area under concentration-time curve (AUC) in PELF were 3.67 ± 0.25 μg/mL and 24.28 ± 2.70 h·μg/mL, higher than those in plasma (0.67 ± 0.01μg/mL and 4.47 ± 0.51 h·μg/mL). The peak time (T_max_) in plasma was 0.23 ± 0.07 h, shorter than that in PELF (1.61 ± 0.15 h). The CO_PD_ in plasma and PELF were 0.125 μg/mL and 0.5 μg/mL, respectively. The CO_CL_ calculated by WindoW approach, nonlinear regression and CART analysis were 0.125∼4 μg/mL, 0.428 μg/mL and 0.56 μg/mL, respectively. The 0.5 μg/mL was selected as eligible CO_CL_. The ECV is much higher than the CO_PD_ and CO_CL_, and the clinical breakpoint based on data in plasma was large different with that of in PELF.

**Conclusions:** Our study firstly established three cutoff values of danofloxacin against *G. parasuis*. It suggested that epidemiological danofloxacin-resistant *G. parasuis* may lead to the ineffective treatment by danofloxacin.

**Importance:** *G. parasuis*, a gram-negative respiratory pathogen, can colonize in the upper respiratory tract in swine and cause Glasser’s disease. As the abuse of antibiotics, antimicrobial resistant *G. parasuis* emerged in different degrees, which brought serious threat to global economy and public health. Danofloxacin in quinolones are one of the best choices for treatment of *G. parasuis* infection, because of their strong bactericidal activity and good absorption into blood and great distribution in the lung. However, the clinical breakpoint (CBP) for danofloxacin against *G. parasuis* had not yet been established by clinical laboratory of standard Institute (CLSI) and European Commission of antimicrobial susceptibility testing (EUCAST). Our study firstly established three cutoff values of danofloxacin against *G. parasuis*. It suggested that epidemiological danofloxacin-resistant *G. parasuis* may lead to the ineffective treatment by danofloxacin.

## 1 Introduction

*Glaesserella parasuis*, a gram-negative respiratory pathogen, can colonize in the upper respiratory tract in swine and cause Glasser’s disease like fibrinous polyserositis, arthritis, meningitis and pneumonia(Oliveira and Pijoan, 2004). The serotype 1, 5, 10, 12, 13 and 14 exhibited higher virulence and pathogenicity(Kielstein and Rapp-Gabrielson, 1992). The serotype 5 and 4 were dominant in China(Cai et al., 2005). As the abuse of antibiotics, antimicrobial resistant *G. parasuis* emerged in different degrees, which brought serious threat to global economy and public health(Nedbalcova et al., 2017).

Quinolones are one of the best choices for treatment of *G. parasuis* infection, because of their strong bactericidal activity and good absorption into blood and great distribution in the lung(Drlica and Zhao, 1997). Danofloxacin, one of the most important fluoroquinolones, has broad spectrum of antimicrobial activity and has been widely used in different animals, like in sheep(Aliabadi et al., 2003), honey(Cherif et al., 2015), rabbits(Fernandez-Varon et al., 2007), turkeys(Haritova et al., 2006), cattle and swine(Mann and Frame, 1992). However, the clinical breakpoint (CBP) for danofloxacin against *G. parasuis* had not yet been established by clinical laboratory of standard Institute (CLSI) and European Commission of antimicrobial susceptibility testing (EUCAST).

The CBP was set on the basis of epidemiological cutoff values (ECV) or wide-type cutoff (CO_WT_), pharmacodynamics (PD) cutoff values (CO_PD_) and clinical cutoff values (CO_CL_)(Toutain et al., 2017). For a given microbial species and antimicrobial agent, the ECVs were the upper bound of the wild-type MIC distribution for organisms without detectable acquired resistance mechanisms and can be calculated by nonlinear regression analysis using ECOFFinder software (Canton et al., 2012; Kronvall, 2010; Turnidge et al., 2006). CO_PD_ considered the pharmacokinetics-pharmacodynamic (PK-PD) parameters of special antimicrobial agent in target animals and determined by Monte Carlo simulation to find the MIC with 90% possibility reaching to the PK-PD target (Rey et al., 2014). CO_CL_ was decided based on the relationship between clinical outcomes and antimicrobial susceptibility using several statistical approaches (Turnidge and Martinez, 2017). The present study was aimed to establish the ECV, CO_PD_ and CO_CL_ values for decision of the final CBP of danofloxacin against *G. parasuis* and evaluation of the efficiency of danofloxacin for treatment of *G. parasuis*.

## 2 Materials and Methods

### 2.1 Strains

From March to May in 2017, a total of 347 *G. parasuis* strains were collected from disease animals. 35 *G. parasuis* strains were isolated from pig lungs provided by Keqian clinical diagnostic center; 8 *G. parasuis* strains were donated by Xiaojuan Xu from State Key Laboratory of Agricultural Microbiology in Huazhong Agricultural University; 204 *G. parasuis* strains were isolated from disease pigs by Peng Zhang in China Agricultural University; and 100 *G. parasuis* strains were stored in National Reference Laboratory of Veterinary Drug Residues. All these strains were isolated from the lungs and pericardium of weak or moribund pigs showing respiratory distress or arthritis in different provinces of China. All bacterial isolates were confirmed by PCR amplification of 16s rRNA(Oliveira et al., 2001). *E*.*coli* (ATCC25922) was used as the quality control (QC) which reserved by National Reference Laboratory of Veterinary Drug Residues.

### 2.2 Animals

78 six-weeks-old healthy crossbred (Duroc × Large × white × Landrace) pigs weighing 20 kg were purchased from Huazhong Agricultural University pig breeding farm. Prior to experiments, pigs were raised 7 days to acclimate. All the animal experiments were approved by the Animal Ethics Committee of Huazhong Agricultural University (hzauch 2014-003) and the Animal Care Center, Hubei Science and Technology Agency in China (SYXK2013–0044). All efforts were used to reduce the pain and adverse effect of the animals.

### 2.3 Establishment of ECV

Susceptibility testing was performed by agar dilution method according to CLSI M07-A9 standard with some modification. A 2 μL *G. parasuis* suspension (10^7^ CFU/mL) was inoculated onto TSA-FCS-NAD agar plates containing two fold dilutions (0.0075……64 μg/mL) of danofloxacin (Dr. Ehrenstorfer Standards, Augsburg, Germany). The MICs were converted to Log_2_MIC, ECV was simulated using ECOFFinder software (Espinel-Ingroff et al., 2018). ECV at 95%, 97.5%, 99%, 99.5% and 99% confidence intervals were simulated.

### 2.4 Establishment of CO_PD_ based on PK-PD modeling

#### 2.4.1 Selection of pathogenic *G. parasuis*

The serotype of 81 strains with MIC same and higher than MIC_90_ were determined by ERIC-PCR using ERIC primer (5’-ATG TAA GCT CCT GGG GAT TCA C-3’ and 5’-AAG TAA GTG ACT GGG GTG AGC G-3’) following previous study (Rafiee et al., 2000;Versalovic et al., 1991). SH 0165 (serotype 5) was positive control.

The 18 strains with serotype 5 were selected to pathogenicity test on mouse. 16 ∼ 20 g healthy Balb/c mice were divided randomly into 19 groups (5 mice/group) with one black control group. 1×10^9^ cfu bacterium was inoculated by abdominal cavity injection, the control group injected with TSB broth. Mice were monitored daily for 7 days post-inoculation (dpi). The pathogenicity of *G. parasuis* was compared based on survival time (Yu et al., 2016).

#### 2.4.2 Pharmacodynamics *in vitro* and *ex-vivo*

The MIC and MBC of *G. parasuis* H80 in broth and pulmonary epithelial lining fluid (PELF) were determined using broth dilution method according to the CLSI M07-A9 standard with some modification.

The *in vitro* and *ex-vivo* killing curve of danofloxacin in broth and in PELF was drawn by monitoring the Colony formed unite (CFU) changes during the incubation of *G. parasuis* H80 under a series concentration of danofloxacin (1/2 to 32 MIC) for continuous time (0, 1, 2, 4, 6, 8, 12 and 24 h).

#### 2.4.3 Animal experiment and sample collection for Pharmacokinetics study

Danofloxacin was administrated to twelve pigs at a single-dose of 2.5 mg/kg b.w. by intramuscular injection. After administration, 2 mL blood samples were obtained at 0, 0.08, 0.17, 0.25, 0.5, 0.75, 1, 1.5, 2, 3, 4, 6, 8, 10, 12, 24, 36, 48 h. Plasma was separated from blood by centrifugation at 3500r/min for 10min.

To collect PELF samples, atropine (0.05 mg/kg) and propofol (9∼15 mg/kg) were given intramuscularly and intravenously 30 min for anesthesia. Standardized Bronchoaveolar Lavage (BAL) was performed as previously described(Giguere et al., 2011; Zhang et al., 2015), with an electronic fiber opticbron choscope (KangmeiGU-180VET) inserted in the right middle lung lobe. The 50mL of normal saline was instilled into the lobe, and was aspirated into a 50mL centrifugal tube. The PELF samples were collected at 0, 0.5, 1, 1.5, 2, 4, 6, 8, 10, 12, 24, 36, 48 h. The PELF was centrifuged at 800 r/min for 10 min.

#### 2.4.4 Quantitation analysis of danofloxacin by HPLC

Quantitation analysis of danofloxacin in PELF and plasma were conducted using high performance liquid chromatography (HPLC). Agent SB-Aq reverse-phase column (250 mm, 4.6 mm i.d., 5 mm; Agilent) was used to perform HPLC at 30°C. The detection wave length was 280nm. The mobile phase consisted of 0.05% phosphoric acid (phase A) and acetonitrile (phase B) with gradient elute. 0.5mLPlasma and 0.5 mL PELF were extracted with 2mL acetonitrile twice.

The urea dilution method was used to determine the volume of PELF as described previously(Conte et al., 2000; Kiem and Schentag, 2008). The concentration of urea in plasma (Urea_PLASMA_) and PELF (Urea_PELF_) were determined by using urea test kit (Urea test kit; Sigma Chemical, St Louis, MO, USA) and the absorbance values measured by using spectrophotometer (Wuhan, China).The final concentration of danofloxacin in PELF (C_PELF_) was derived from the following equation: 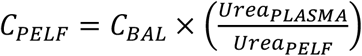, C_BAL_ was diluted concentration of danofloxacin in PELF determined by HPLC method.

#### 2.4.5 Pharmacokinetics-pharmacodynamics modeling

PK-PD parameters were analyzed with a two-compartment model by Winnonlin v.5.2.1. According to the *ex-vivo* time-killing curve, AUC_24_/MIC (AUIC) of danofloxacin under different concentrations was calculated by Sigmoid E_max_ model 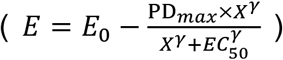, E is the summary PD endpoint, and E_0_ is the effect representing the value of the PD endpoint without drug treatment (i.e., the value of the summary endpoint when the PK/PD index is 0). X is one of the three PK/PD indices as defined above, PD_max_ is the maximum effect (in relation to E_0_) indicated by the plateau where increased exposures result in no further kill. EX_50_ is the magnitude of X that is needed to achieve 50% of PD_max_, and γ is the sigmoidicity factor. The PD target was determined with *Sigmoid E*_*max*_ equation (Mouton, 2002; Xiao et al., 2015). The dosage regimen was derived from the concentration-dependent dosage equation 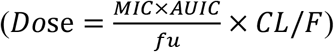(Potter et al., 2013; Sidhu et al., 2014; Toutain et al., 2002). In the equation, the CL was the plasma (total) clearance in days, fu was the free fraction of the drug in plasma (from 0 to 1), and F% was the bioavailability factor (from 0 to 1).

#### 2.4.6 Monte Carlo Simulation to set up CO_PD_

Crystal Ball v7.2.2 was used to perform Monte Carlo simulation. The distribution of pharmacokinetic parameter AUC_24_ was assumed to be log-normal. A total of 10000 subjects were simulated. The PD target was selected to calculate the probability of target attainment (PTA). CO_PD_ was defined as the MIC at which the PTA was ≥90%.

### 2.5 Clinical trial and Establishment of CO_CL_

#### 2.5.1 Infection model and Clinical trials

66 healthy weaned piglets (about 20 kg) were divided into 11 groups, 5 groups were experimental group, 5 groups were negative control group, and 1 group was blank control group, 6 piglets in each group. The 5 experimental groups and 5 negative control groups were challenged with 5 representative strains H42, H80, H12, H83 and H17 by intranasal inoculation of 1×10^10^ CFU bacterial suspension twice a day. The blank control group was inoculated with blank TSB broth. The dosage regimens were recommended by PK-PD therapeutic dosage regimen. After challenging, these pigs were monitored daily for two weeks.

#### 2.5.2 Statistical analysis for establishment of CO_CL_

The probability of cure (POC) was calculated based on the clinical outcomes and bacteriological prognosis. Clinical outcomes included treatment success and failure, and each MIC should have a corresponding clinical outcome. Bacteriological prognosis was to determine the presence or eradication of the bacteria after administration. The data were analyzed by three different analysis methods.

The “WindoW” approach (Turnidge and Martinez, 2017) included two parameters: “MaxDiff” and “CAR”. “MaxDiff (the method of maximum difference, MaxDiff)” represents the difference between higher and lower POC at a certain MIC. “CAR” was based on the clinical outcome and the corresponding MIC distribution. “CAR” could not be set as the lowest MIC or the highest MIC, if CAR was gradually increasing with MIC, then the CAR should choose the second small CAR.

Nonlinear regression analysis was a new method based on the formula between EUCAST proposed POC with MIC. Log_2_MIC was independent variable, the POC was dependent variable. The model with highest correlation coefficient was selected to simulate its CO_CL_.

The classification and regression tree (CART) model (Salford Predictive Modeler software) was also used for establishment of CO_CL_. MIC was used as the predictive variable and POC was the target variable. The Gini coefficient minimization criterion was used to select the MIC node automatically.

## 3 RESULTS

### 3.1 ECV for danofloxacin against *G. parasuis*

The MIC distribution for danofloxacin against *G. parasuis* was shown in Figure 1. The MIC of danofloxacin was ranged from 0.008 to 64 μg/mL. As shown in figure 1, the MIC distribution was as follows: 0.008 µg/mL (2.88%), 0.015 µg/mL (1.15%), 0.03 µg/mL (5.19%), 0.06 µg/mL (6.34%), 0.125 µg/mL (7.20%), 0.25 µg/mL (5.48%), 0.5 µg/mL (2.88%), 1 µg/mL (8.36%), 2 µg/mL (27.09%), 4 µg/mL (19.60%), and 8 µg/mL (8.65%), 16 µg/mL (4.33%), 32 µg/mL (0.58%), 64 µg/mL (0.29%). The MIC_50_ and MIC_90_ were 2 μg/mL and 8 μg/mL, respectively.

**Figure 1.**
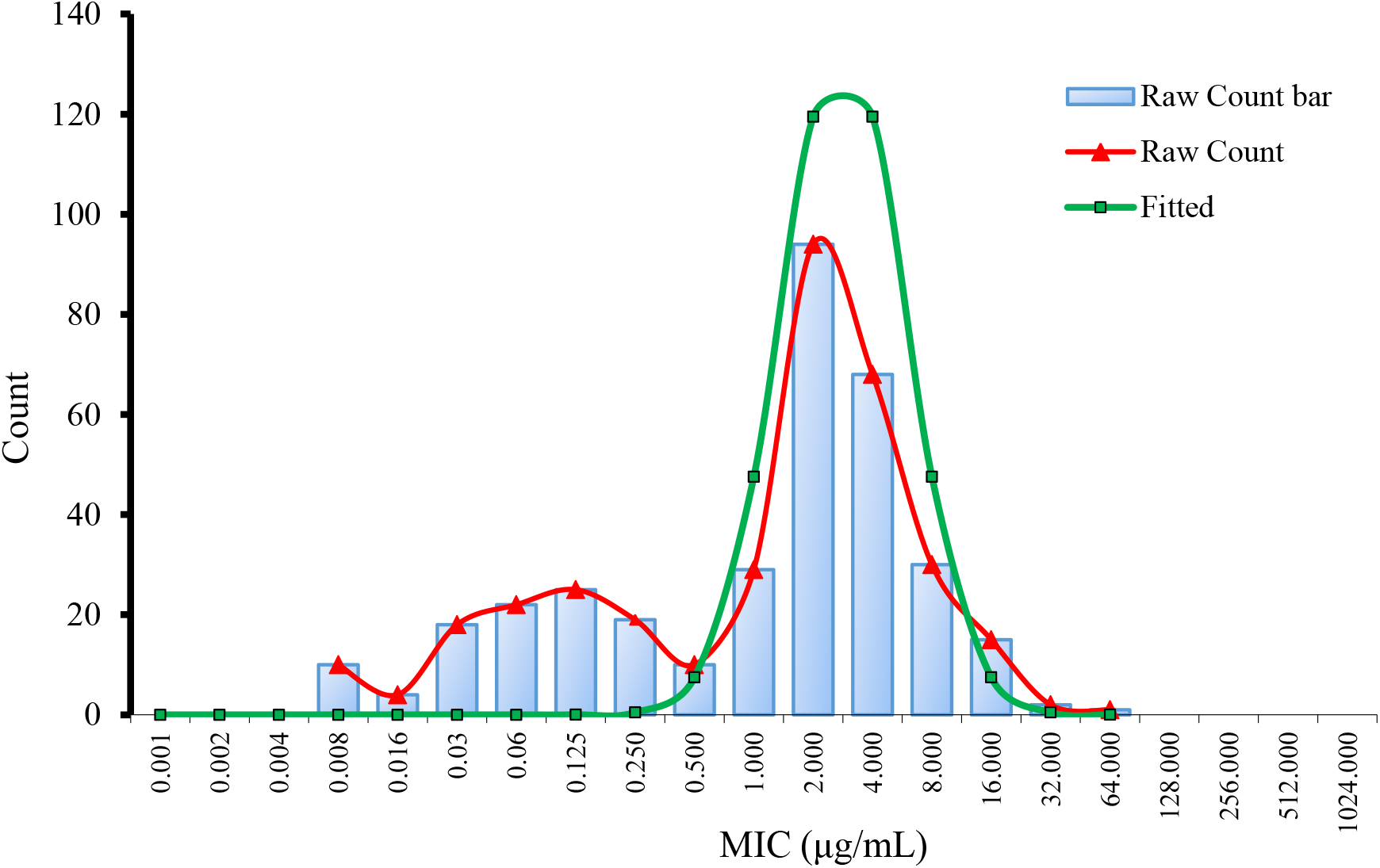
Nonlinear regression of MIC distribution for danofloxacin against *G. parasuis*

Using ECOFFinder software, the fitted MIC distribution of danofloxacin against *G. parasuis* was shown in Figure 1. The ECV at 95%, 97.5%, 99%, 99.5% and 99.9% confidence intervals were 8, 8, 16,16 and 32 μg/mL, respectively.

### 3.2 CO_PD_ for danofloxacin against *G. parasuis*

#### 3.2.1 Pathogenic *G. parasuis*

18 strains with serotype 5 were selected from ERIC-PCR amplificatioan and pathogenicity test in mice and five strains (H42, H80, H12, H83 and H17) showed highest pathogenicity and exhibited different MIC. The strain H80 with MIC close to MIC_50_ was selected for PK-PD study. The 5 respective strains H42 (MIC=16 µg/mL), H80 (MIC=4 µg/mL), H12 (MIC=1 µg/mL), H83 (MIC=0.125 µg/mL) and H17 (MIC=0.015 µg/mL) were selected for clinical trial.

#### 3.2.2 Pharmacodynamics of danofloxacin against *G. parasuis*

The MIC of danofloxacin in broth and pulmonary epithelial lining fluid (PELF) were 4 μg/mL and 2 μg/mL, respectively. The MBC in broth and PELF were 8 μg/mL and 4 μg/mL, respectively. The antibacterial activity of danofloxacin in PELF is stronger than that of in broth. The PELF may contain certain antibodies or immunological factors or other chemicals which can enhance the antibacterial activity of danofloxacin.

As displayed in Figure 2A/B, the *in vitro* and *ex-vivo* bactericidal effect of danofloxacin against *G. parasuis* was similar. The lower concentrations (≤ MIC) of danofloxacin exhibited similar antibacterial activity to *G. parasuis*. However, when danofloxacin concentrations were higher than MIC, the inhibitory efficiency gradually strengthened following the increased drug concentration. The time killing curve showed that activity of danofloxacin against *G. parasuis* was concentration-dependent. The Aera Under Curve/ Minimum Inhibitory Concentration (AUC/MIC) was selected as PK-PD parameter.

**Figure 2.**
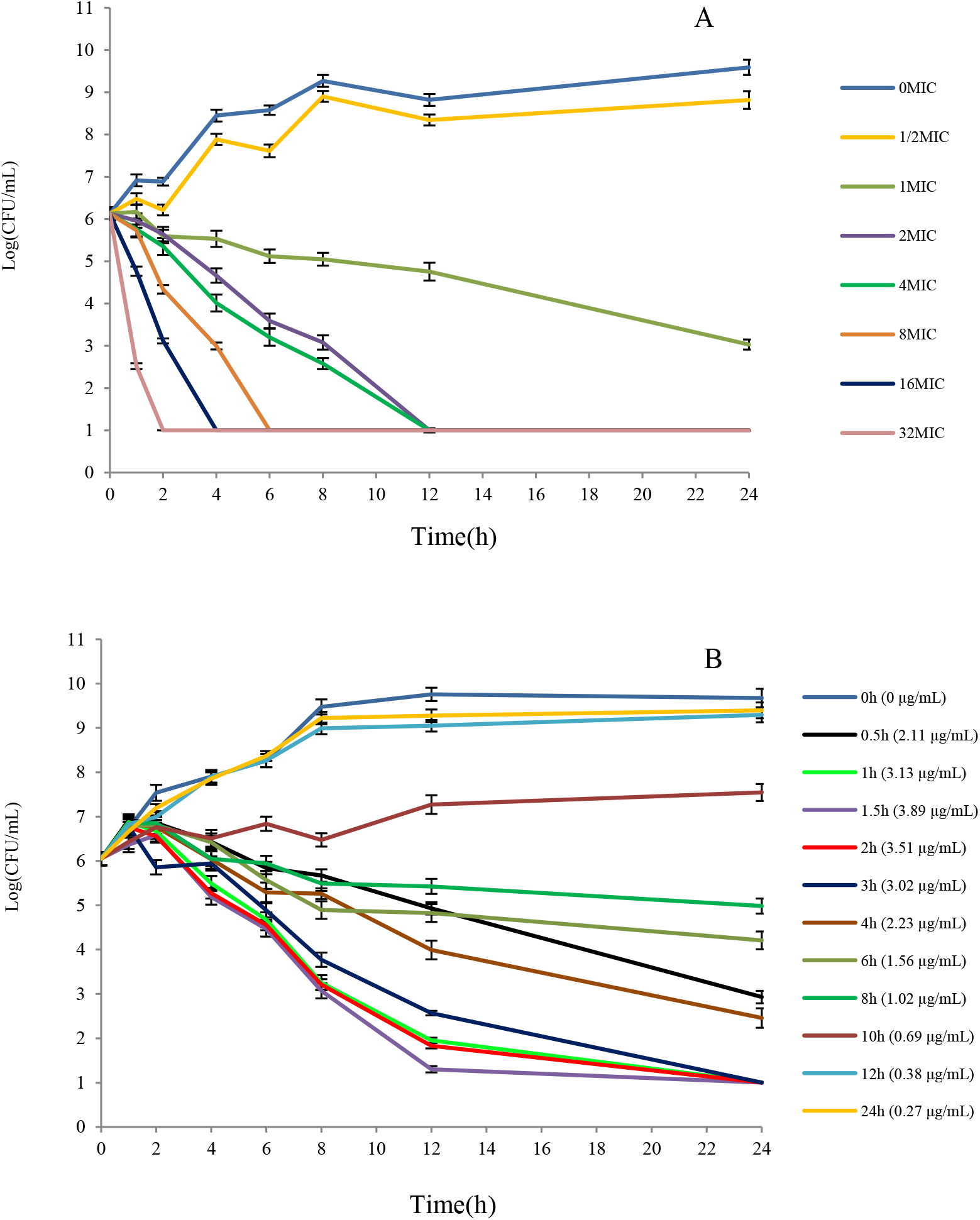
A/B The killing curve of *G. parasuis* in PELF and plasma Note: Figure A was the killing curve of *G. parasuis* in TSB broth, Figure B was killing curve of *G. parasuis* in PELF.

#### 3.2.3 Sensitivity and accuracy of HPLC method for determination of danofloxacin

The limit of determination (LOD) was 0.01 μg/mL and the limit of quantification (LOQ) was 0.025 μg/mL in PELF. The LOD was 0.02 μg/mL and the LOQ was 0.05 μg/mL in plasma. Standard curves were linear from 0.05 μg/mL to 5 μg/mL in plasma (R^2^ = 0.9994) and 0.025 μg/mL to 2.5 μg/mL in PELF (R^2^ = 0.9996). The inter-day variation for determination in plasma and PELF ranged from 1.94% to 2.37% and 1.36% to 2.71%, respectively. The recovery of danofloxacin in plasma and PELF ranged from 90.79±2.15 to 94.36±1.83 and 91.91±2.49 to 95.73±1.30, respectively.

#### 3.2.4 PK characteristics of danofloxacin in plasma and PELF

The concentration-time curves in plasma and PELF after administration of danofloxacin at a single dose of 2.5 mg/kg b.w. were shown in Figure 3. Significant differences were observed between drug concentrations in plasma and in PELF.

**Figure 3.**
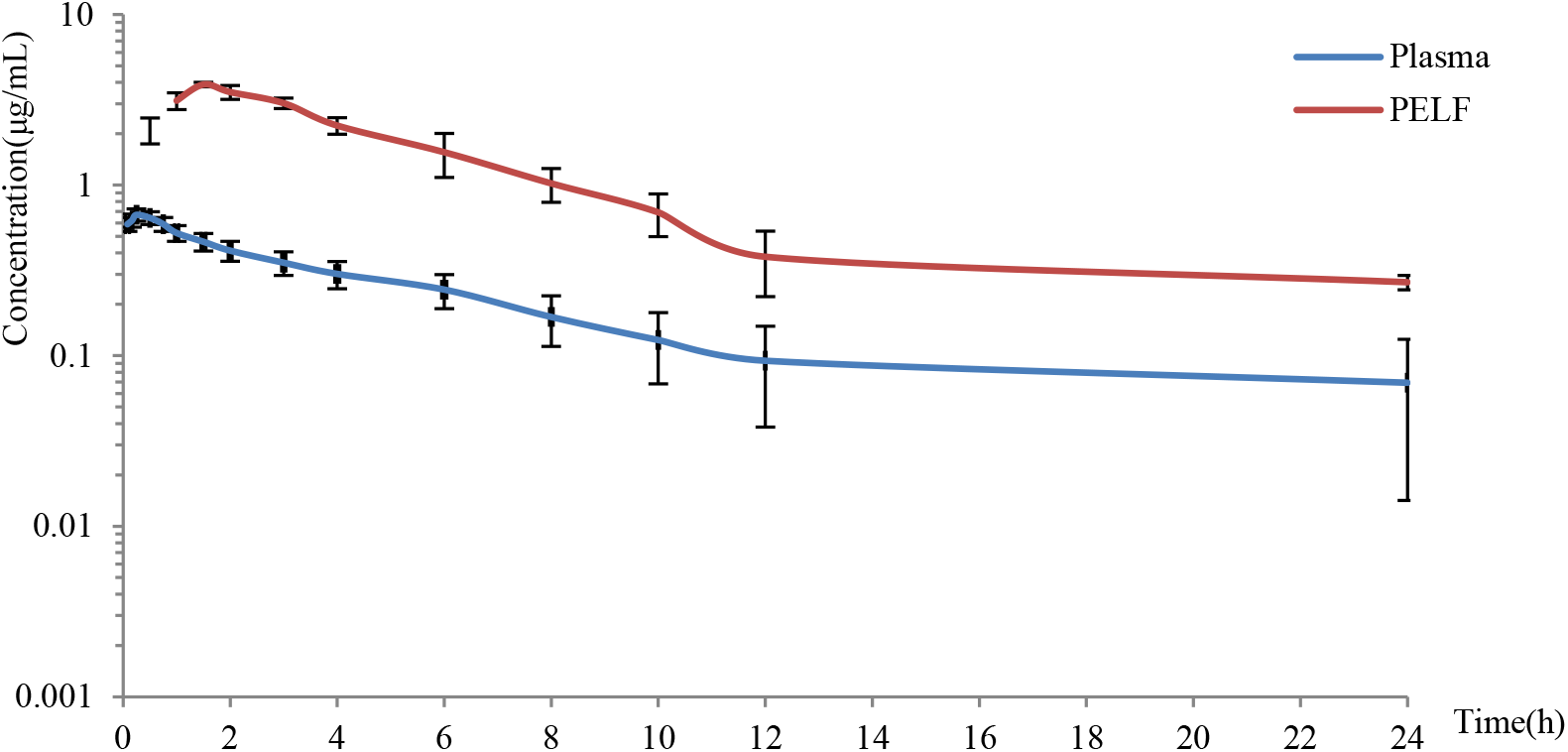
Mean concentration versus time curves for danofloxacin in PELF and plasma

The simulated pharmacokinetic parameters in plasma and PELF were shown in Table 1. In plasma, the peak time (T_max_) was 0.23 ± 0.07 h, the peak concentration (C_max_) was 0.67 ± 0.01 μg/mL, the area under the concentration-time curve (AUC) was 4.47 ± 0.51 h·μg/mL; in PELF, T_max_ was 1.61 ± 0.15 h, C_max_ was 3.67 ± 0.25 μg/mL, AUC was 24.28 ± 2.70 h·μg/mL.

**Table 1.** PK parameters of danofloxacin in plasma and PELF (n=6)

| Parameters | Unit | Plasma | PELF |
| --- | --- | --- | --- |
| A | μg/mL | 0.43±0.16 | 6.50±2.21 |
| B | μg/mL | 0.37±0.18 | 0.54±0.40 |
| α | 1/h | 0.40±0.13 | 0.29±0.04 |
| β | 1/h | 0.14±0.02 | 0.06±0.02 |
| T<sub>1/2α</sub> | h | 1.78±0.76 | 2.39±0.3 |
| T<sub>1/2β</sub> | h | 4.96±0.47 | 10.46±0.76 |
| T<sub>max</sub> | h | 0.23±0.07 | 1.61±0.15 |
| AUC<sub>24</sub> | h·μg/mL | 4.47±0.51 | 24.28±2.70 |
| C<sub>max</sub> | μg/mL | 0.67±0.01 | 3.67±0.25 |
| CL/F | mL/h/kg | 571.49±53.02 | 89.98±9.7 |
| Vd/F | mL/kg | 435.04±45.43 | 3531.73±49.12 |

Combined with the killing curve in PELF, the PD target (AUIC in *ex-vivo*) under different efficiency was calculated by Sigmoid E_max_ equation simulation (Table 2). The values of AUIC (h) at E = 0, −3 and −4 (bacteriostasis, bactericidal and eradication) were 12.73, 28.26 and 44.38, respectively.

**Table 2.**
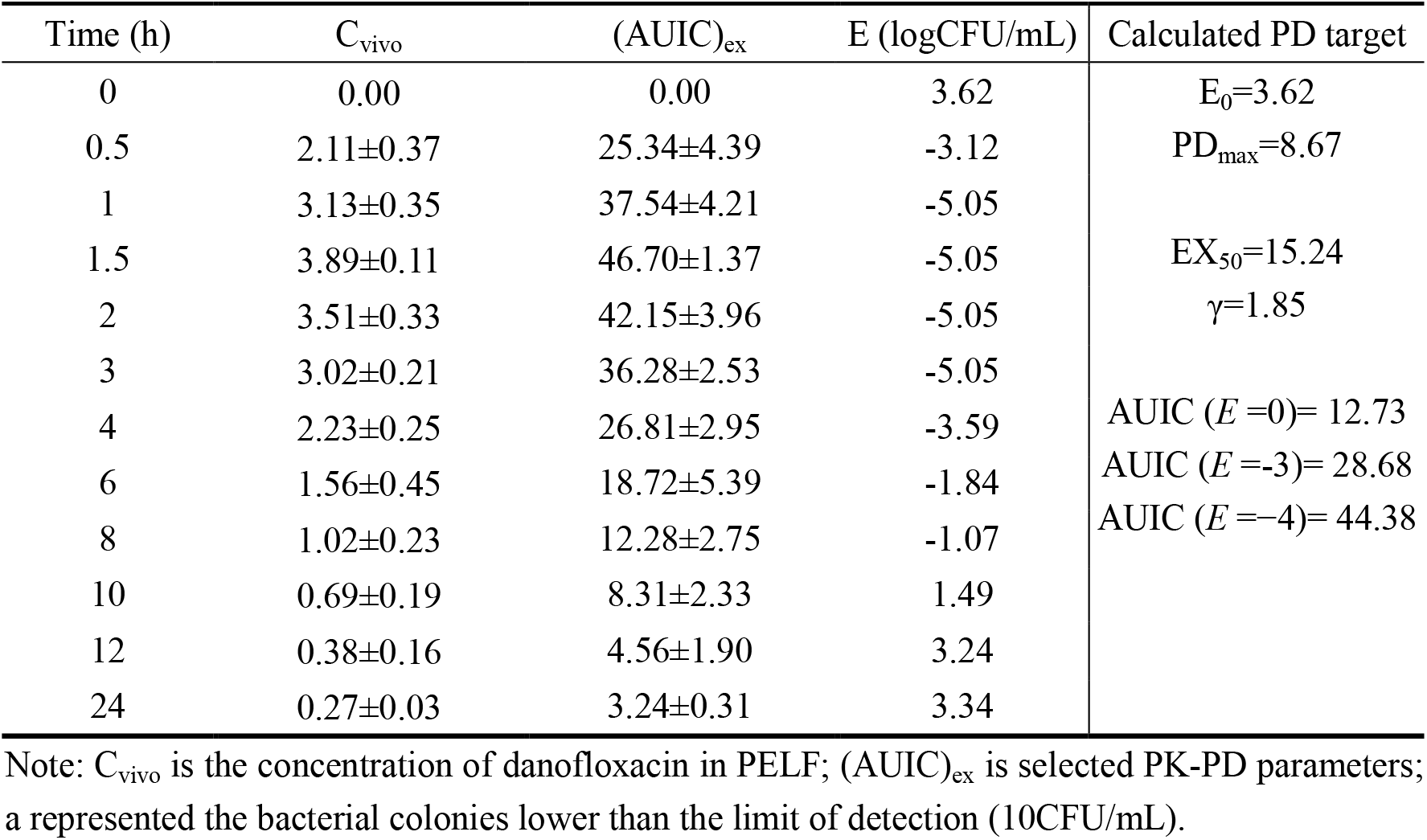
PD target of danofloxacin against *G. parasuis*

#### 3.2.5 Monte Carlo Simulation and CO_PD_

According to the AUC (24.28 ± 2.70 h·μg/mL) and PD target (12.73, 28.26, 44.38) in PELF, the possibility of target achievement (PTA) at different MIC was simulated by the Monte Carlo analysis (Table 3). When the PTA in PELF was upon 90%, the CO_PD_ (E=0, −3, −4) for danofloxacin against *G. parasuis* in PELF was 1 μg/mL, 0.5 μg/mL, 0.25 μg/mL, respectively.

**Table 3.**
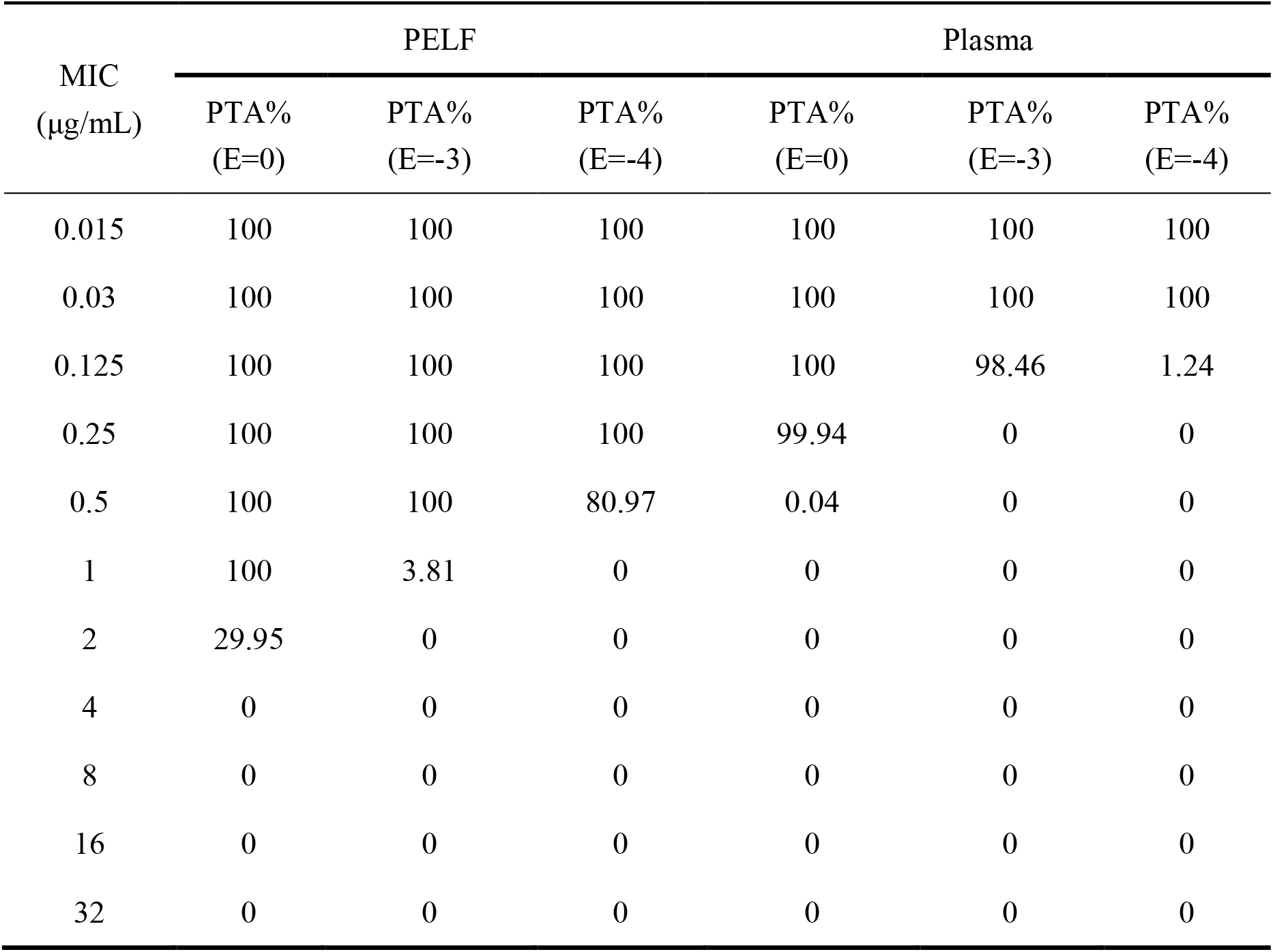
PTA of danofloxacin against *G. parasuis* at different MIC in PELF and plasma

| MIC<br>(µg/mL) | PELF |  |  | Plasma |  |  |
| --- | --- | --- | --- | --- | --- | --- |
|  | PTA%<br>(E=0) | PTA%<br>(E=-3) | PTA%<br>(E=-4) | PTA%<br>(E=0) | PTA%<br>(E=-3) | PTA%<br>(E=-4) |
| 0.015 | 100 | 100 | 100 | 100 | 100 | 100 |
| 0.03 | 100 | 100 | 100 | 100 | 100 | 100 |
| 0.125 | 100 | 100 | 100 | 100 | 98.46 | 1.24 |
| 0.25 | 100 | 100 | 100 | 99.94 | 0 | 0 |
| 0.5 | 100 | 100 | 80.97 | 0.04 | 0 | 0 |
| 1 | 100 | 3.81 | 0 | 0 | 0 | 0 |
| 2 | 29.95 | 0 | 0 | 0 | 0 | 0 |
| 4 | 0 | 0 | 0 | 0 | 0 | 0 |
| 8 | 0 | 0 | 0 | 0 | 0 | 0 |
| 16 | 0 | 0 | 0 | 0 | 0 | 0 |
| 32 | 0 | 0 | 0 | 0 | 0 | 0 |

According to the AUC (4.47 ± 0.51 h·μg/mL) and PD target (12.73, 28.26, 44.38) in plasma, the PTA at different MIC was simulated by the Monte Carlo analysis (Table 3). When the PTA in plasma was upon 90%, the CO_PD_ (E=0, −3, −4) for danofloxacin against *G. parasuis* in plasma was 0.25 μg/mL, 0.125 μg/mL, 0.03 μg/mL, respectively.

### 3.3 CO_CL_ of danofloxacin against *G. parasuis*

The dosage under different efficiency (bacteriostasis, bactericidal and eradication) were 4.58 mg/kg, 10.32 mg/kg and 15.97 mg/kg. The given dosages were simulated by Mlxplore software. The modified dosage regimen was 12.49 mg/kg danofloxacin twice a day. Three methods were used to obtain CO_CL_ according to the relationship between POC and MIC distribution (Table 4).

**Table 4.** POC and “WindoW” for danofloxacin against *G. parasuis* at different MIC

| Strain number | Strain group | MIC (µg/mL) | Success (%) | Eradication (%) | POC (%) | MaxDiff | CAR |
| --- | --- | --- | --- | --- | --- | --- | --- |
| H42 | Test | 16 | 67.7 | 67.7 | 67.7 | 0 | 0.70 |
|  | Controll |  | 16.7 | 0 | 0 |  |  |
| H80 | Test | 4 | 67.7 | 83.3 | 67.7 | 0 | 0.79 |
|  | Controll |  | 33.3 | 16.7 | 33.3 |  |  |
| H12 | Test | 1 | 83.3 | 83.3 | 83.3 | 0.167 | 0.93 |
|  | Controll |  | 33.3 | 16.7 | 33.3 |  |  |
| H83 | Test | 0.125 | 100 | 100 | 100 | 0.28 | 1 |
|  | Controll |  | 33.3 | 16.7 | 16.7 |  |  |
| H17 | Test | 0.015 | 100 | 100 | 100 | 0.21 | 1 |
|  | Controll |  | 50 | 33.3 | 33.3 |  |  |

Following “WindoW” method, the parameters of MaxDiff (0.28) and CAR (0.78) was corresponding with the MIC of 0.125 μg/mL and 4 μg/mL, respectively. The selection window for CO_CL_ was therefore ranged from 0.125 μg/mL to 4 μg/mL.The nonlinear regression model was set up as *y* = 80.989 − 7.271*x* + 0.271*x*^2^ + 0.16*x*^3^ with the correlation coefficient of 0.996. When POC was 90%, the recommended CO_CL_ (MIC) was less than 0.428 μg/mL. The CART regression tree indicated that the CO_CL_ was less than 0.56 μg/mL. Combined with the above three results, the CO_CL_ of danofloxacin against *G. parasuis* was selected as 0.25 μg/mL.

## 4 Discussion

*G. parasuis is an important pathogen for* respiratory infection in swine. Antimicrobial treatment is the best way to control this pathogen due the vaccine deficiency. However, antimicrobial resistance in *G. parasuis* had been found in Germany(Aarestrup et al., 2004), United Kingdom, Spain(de la Fuente et al., 2007) and China(Wang et al., 2017; Xu et al., 2011; Zhou et al., 2010). To rational use of antimicrobial agents to control *G. parasuis*, some studies were carried out to establish the ECVs and/or CO_PD_ of marbofloxacin, cefquinome and tilmicosin against *G. parasuis* (Sun et al., 2015; Xiao et al., 2015; Zhang et al., 2016). The efficiency of danofloxacin on *Actinobacillus pleuropneumoniae* (Lauritzen et al., 2003), *Pasteurella multocida* (Zeng et al., 2011), and *Mannheimia haemolytica* (Fajt et al., 2004) was very good. However, the clinical breakpoint of danofloxacin against *G. parasuis* had not yet been established.

Statistical analysis had been widely used for determination of ECVs. Turnidge(Turnidge et al., 2006) recommend to use nonlinear regression to analyze the obtained MIC data and determined the ECVs of various drugs. Kronvall (Kronvall et al., 2006) used NRI (Normalized Resistance Interpretation) method to analyzed MIC data obtained by E test for establishment of ECVs. European Commission of Antimicrobial Susceptibility Testing (EUCAST) recommended ECOFFinder software on the basis of Turnidge’s nolinear regression (Ismail et al., 2018). Van Vliet(Van Vliet et al., 2017) used NRI and ECOFFinder analysis method to analyze wild type cutoff values of ampicillin, florfenicol, gentamicin and enrofloxacin. In our study, the ECV of danofloxacin determined by nonlinear regression analysis was same with that simulated by ECOFFinder software, suggesting that ECOFFinder software was a convenient tool for establishment of ECVs. In the present study, the MIC distribution of danofloxacin against *G. parasuis* appeared three peaks (0.008 μg/mL, 0.125 μg/mL and 2 μg/mL), suggesting that some *G. parasuis* isolates may be resistant to danofloxacin. Zhang et al. (Zhang et al., 2013) examined the resistance of 138 *G. parasuis* strains against fluoroquinolone drugs and showed that 60.1% isolates was resistant to enrofloxacin and 5.8% isolates were resistant to levofloxacin. It suggested that *G. parasuis* may be also resistant to danofloxacin due to the cross resistance between fluoroquinolone drugs.

The CO_PD_ was established based on pharmacokinetic data, MIC distribution and PK-PD target. Our present study establish the CO_PD_ based on the PK data from healthy animals because of the stability and repeatability of healthy animal model. Considering the drug concentrations in the target sites were directly correlated with clinical efficacy, the PK data both in plasma and in PELF were included into our study(Barbour et al., 2010). Similar with previous studies, our results indicated that the concentration and AUC of danofloxacin in PELF(in lung) was 4∼7 times higher than that in plasma (Mann and Frame, 1992). The CO_PD_ of danofloxacin in PELF was subsequently higher than the CO_PD_ in plasma, indicating that the CO_PD_ was different between in the target issue and in plasma. As danofloxacin can be accumulated at the infection site (lung), the CO_PD_ in plasma may not represent the critical value of the target tissue. It was of great significance to establish the CO_PD_ in target tissue and plasma simultaneously.

Previously, Rowan’s study exhibited good clinical outcome of danofloxacin in the treatment of respiratory disease caused by *Haemophilus somnus* and *Pasteurella multocida* in European cattle (Rowan et al., 2004). The clinical data in our study also showed good clinical outcome of danofloxacin in the treatment of *G. parasuis* in pigs because the success rate for treatment of *G. parasuis* with MIC of 1μg/mL was still as high as 83.33%. The CO_CL_ was established based on relationship between MIC and POC under modified therapeutic dosage. Since there was no standard approach for establishment of CO_CL_, the CO_CL_ in the present study was established by the combination of the three approaches which included the “WindoW” approach (Turnidge and Martinez, 2017), the nonlinear regression (Toutain, 2015), and the CART analysis (Esterly et al., 2012; Zheng et al., 2016). The “WindoW” approach was recommended by CLSI (Turnidge and Martinez, 2017). The nonliner regression with the formula of *POC=1/ (1+e*^*-a+bf (MIC)*^) was proposed by VetCAST to calculate the relation between the dependent variable of POC and the independent variable of MIC (Toutain, 2015). The CART method was previously used to develop clinical breakpoints of cefepime (Bhat et al., 2007) and this method was recommended by Dr Cuesta (Cuesta et al., 2010) and Dr.Toutain (Toutain et al., 2017) because the CART obtained the best statistical results when it was compared with other four supervised classifiers (J48, the OneR decision rule, the naïve Bayes classifier, and simple logistic regression).

The large difference was observed between three cutoff values with ECV higher than CO_PD_ and CO_CL_. In previous studies, Sweeney’s datashowed that the MIC breakpoint of danofloxacin on *Mannheimia haemolytica* and *Pasteurella multocida* were 1μg/mL(Sweeney et al., 2017), while Yang’s data showed that the epidemiologic cutoff value of danofloxacin *Escherichia coli* was 8 μg/mL(Yang et al., 2019), which was in accordance with our study. The difference of ECV between different studies may due to the epidemiological characteristic of different bacterial in different geography. Additionally, previous data showed that some of G.*parasuis* isolates exhibited decreased sensitivity to fluoroquinolones (Guo et al., 2011). Two peak of MIC distribution in the present data also suggested that some G.*parasuis* isolates may be resistant to danofloxacin. The higher MIC of the resistant isolates may contribute to the higher ECV value and further studies may need to confirm the relationship between MIC phenotype and resistance genotype.

## 5 Conclusions

This study firstly established the ECV (8μg/mL) at 95% confidence intervals, CO_PD_ in PELF (0.5 μg/mL), CO_PD_ in plasma (0.125 μg/mL) and CO_CL_ (0.25 μg/mL) of danofloxacin against *G. parasuis*. Based on CLSI decision tree, final CBP in plasma and PELF was 0.25μg/mL and 8 μg/mL, respectively. The ECV value was higher than CO_PD_ and CO_CL_, indicating that some *G. parasuis* isolates may be resistance to danofloxacin.

## Acknowledgments

We appreciated all the participants for their participation and trust and thank Xiaojuan Xu and Hongyan Yu from State Key Laboratory of Agricultural Microbiology in Huazhong Agricultural University for freely providing *G. parasuis* isolates and epidemic materials.

## Transparency declarations

All authors: none to declare.

## Supplementary materials

**Figure 1.**
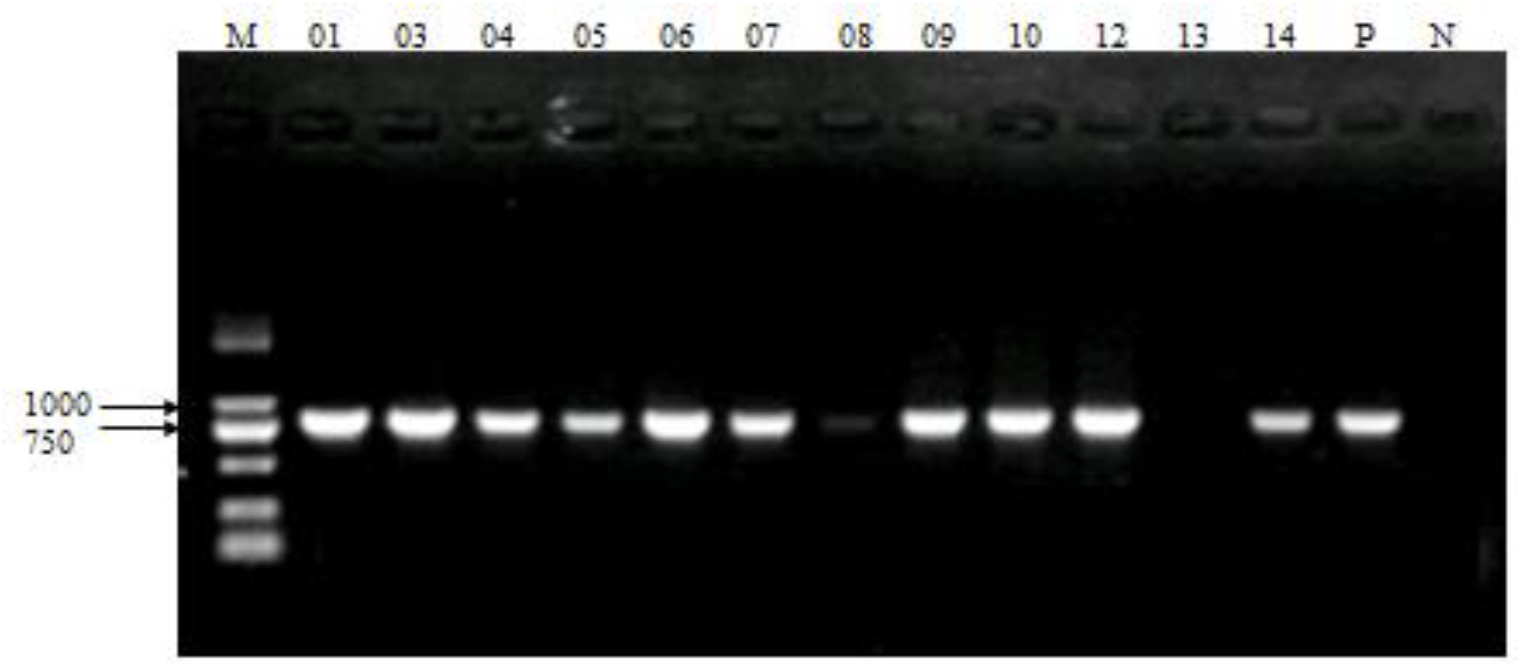
Amplification of *G. parasuis* 16S rRNA with PCR Lane M: DL-2000 DNA Marker; Lane P: positive control; Lane N: Negative control; Lane2-13: Samples to be amplificated.

**Figure 2.**
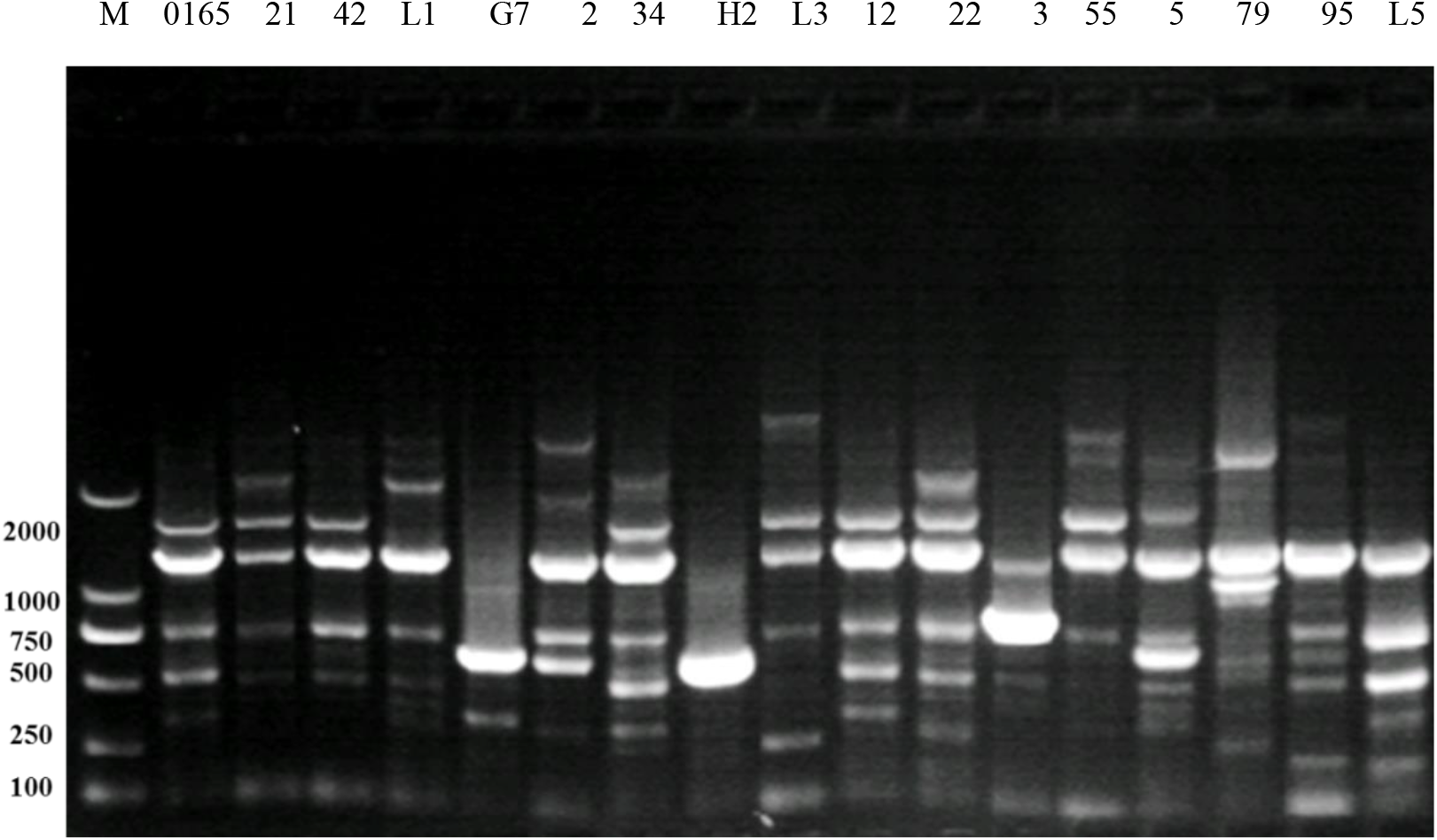
Results of ERIC-PCR for *G. parasuis* Lane M: DL-2000 DNA Marker; Lane 2: SH0165 strain; Lane 17: Negative control; Lane 3-16: Samples to be amplificated

**Figure 3.**
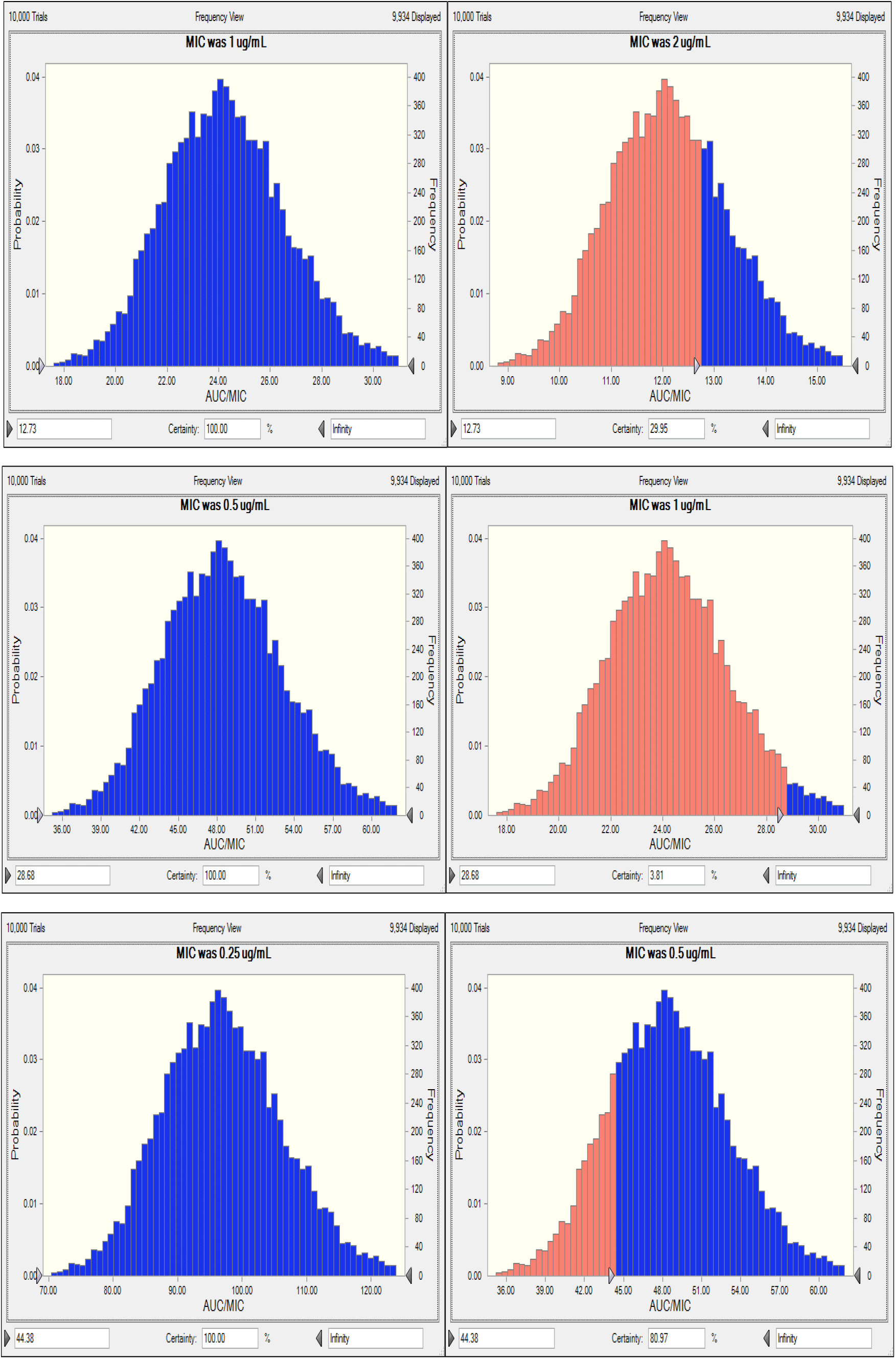
PTA of danofloxacin against *G. parasuis* in PELF

**Figure 4.**
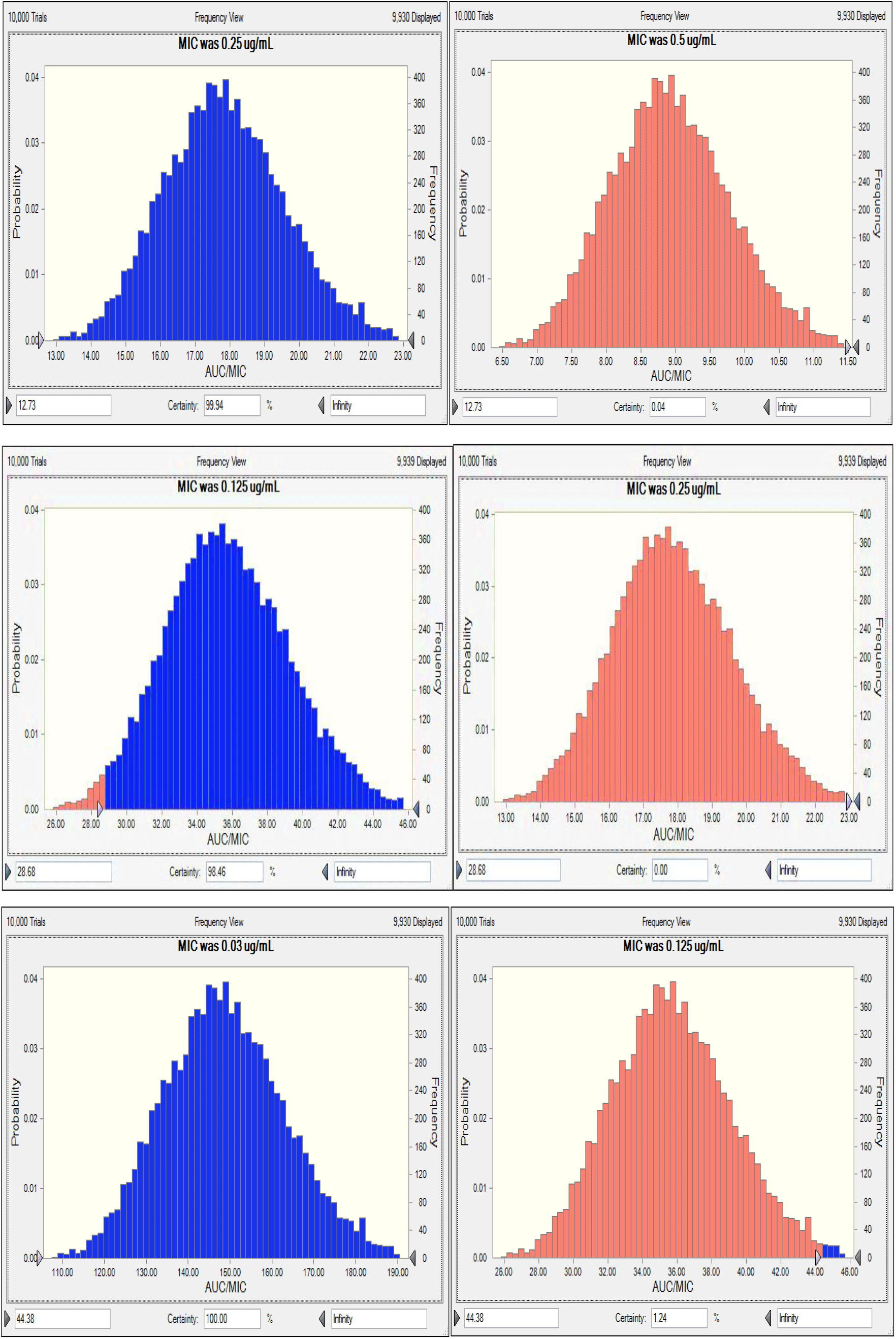
PTA of danofloxacin against *G. parasuis* in plasma

**Figure 5.**
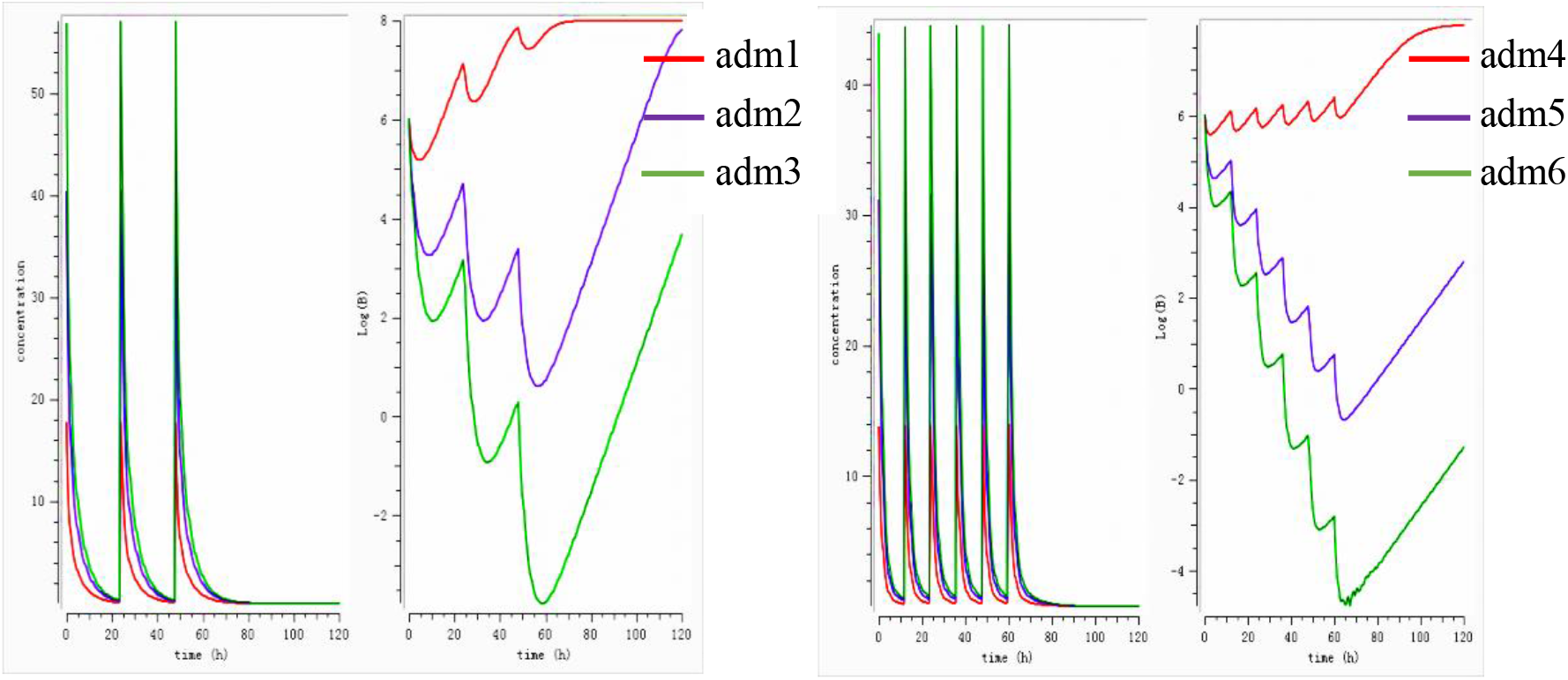
Forecast growth of *G. parasuis* at different dosage regimens Note: adm 1: prevent dosage: 4.58 mg/kg once daily; adm 2: therapeutic dosage: 10.32 mg/kg once daily; adm 3: eradicate dosage: 15.97 mg/kg once daily; adm 4: prevent dosage: 4.58 mg/kg twice daily; adm 5: therapeutic dosage: 10.32 mg/kg twice daily; adm 6: eradicate dosage: 15.97 mg/kg twice daily.

**Figure 6.**
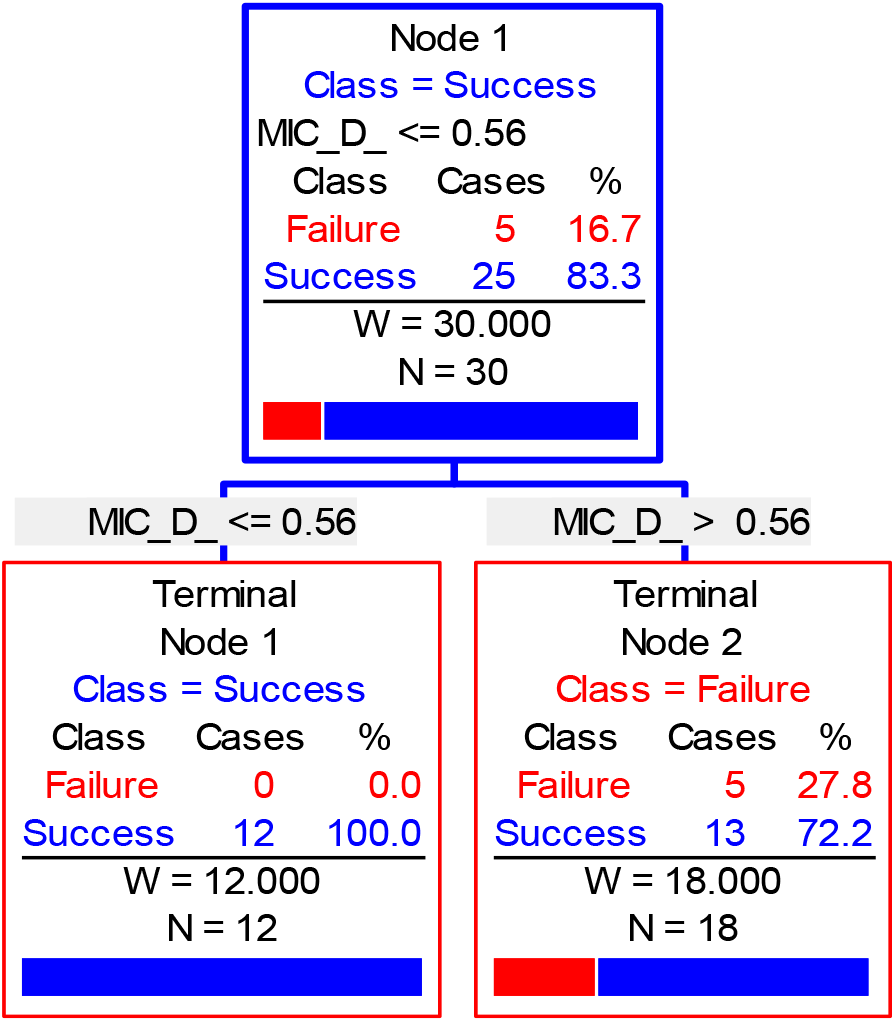
CART tree showing values of clinical outcome

**Figure 7.**
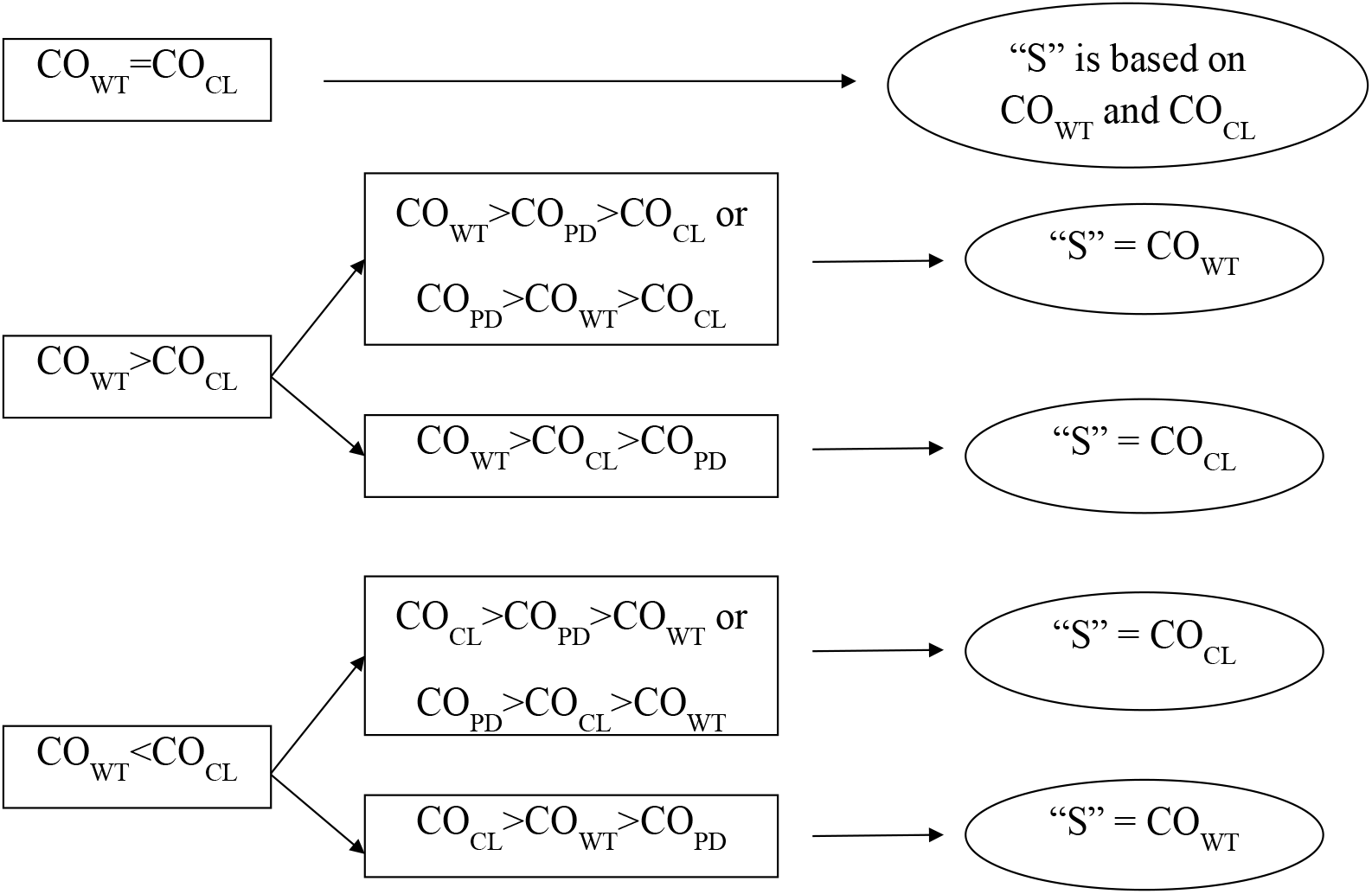
Susceptibility breakpoint decision tree

**Table 1.**
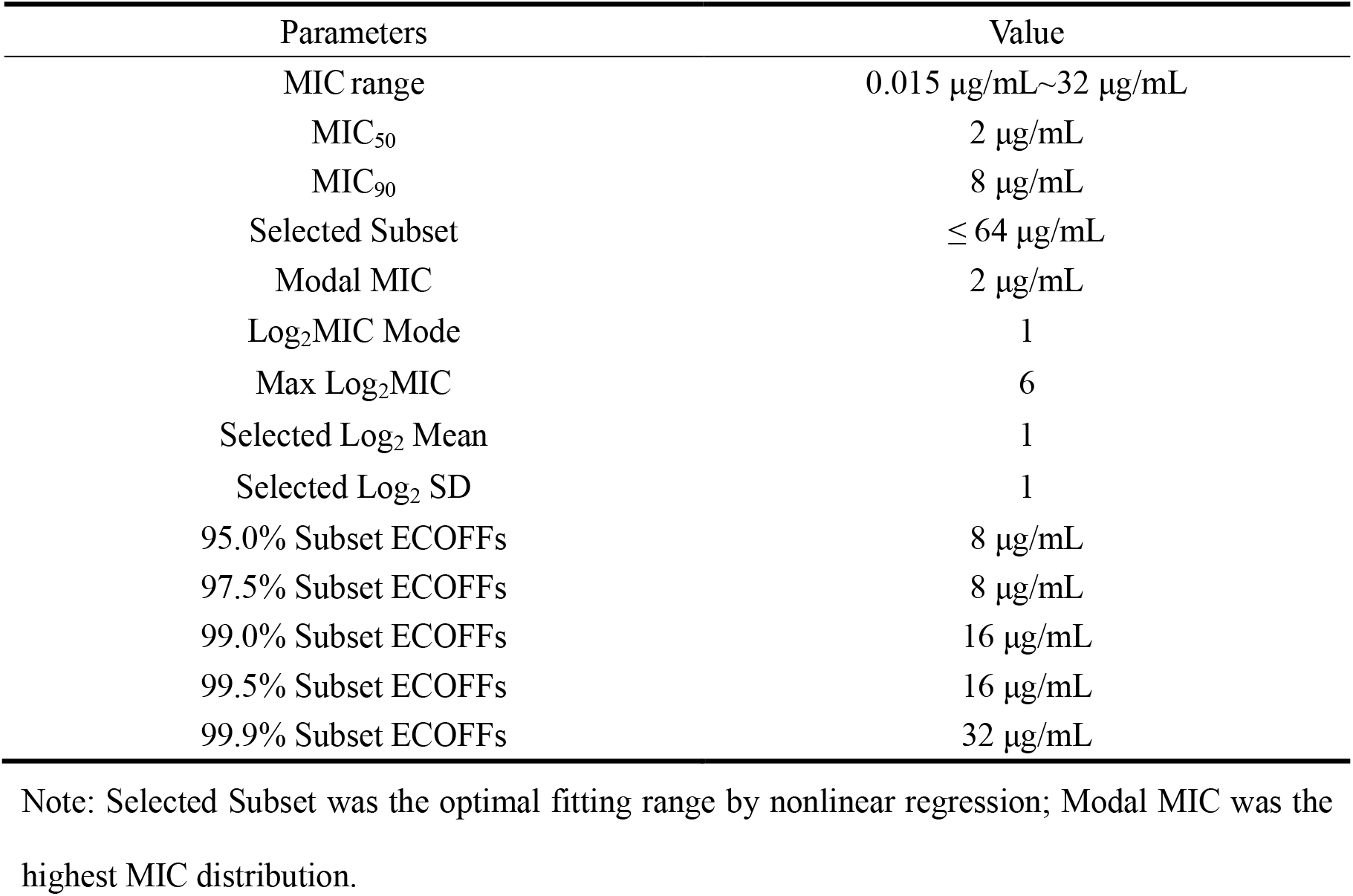
Epidemiological MIC for danofloxacin against *G. parasuis*

| Parameters | Value |
| --- | --- |
| MIC range | 0.015 µg/mL~32 µg/mL |
| MIC<sub>50</sub> | 2 µg/mL |
| MIC<sub>90</sub> | 8 µg/mL |
| Selected Subset | ≤ 64 µg/mL |
| Modal MIC | 2 µg/mL |
| Log<sub>2</sub>MIC Mode | 1 |
| Max Log<sub>2</sub>MIC | 6 |
| Selected Log<sub>2</sub> Mean | 1 |
| Selected Log<sub>2</sub> SD | 1 |
| 95.0% Subset ECOFFs | 8 µg/mL |
| 97.5% Subset ECOFFs | 8 µg/mL |
| 99.0% Subset ECOFFs | 16 µg/mL |
| 99.5% Subset ECOFFs | 16 µg/mL |
| 99.9% Subset ECOFFs | 32 µg/mL |

Note: Selected Subset was the optimal fitting range by nonlinear regression; Modal MIC was the highest MIC distribution.

**Table 2.**
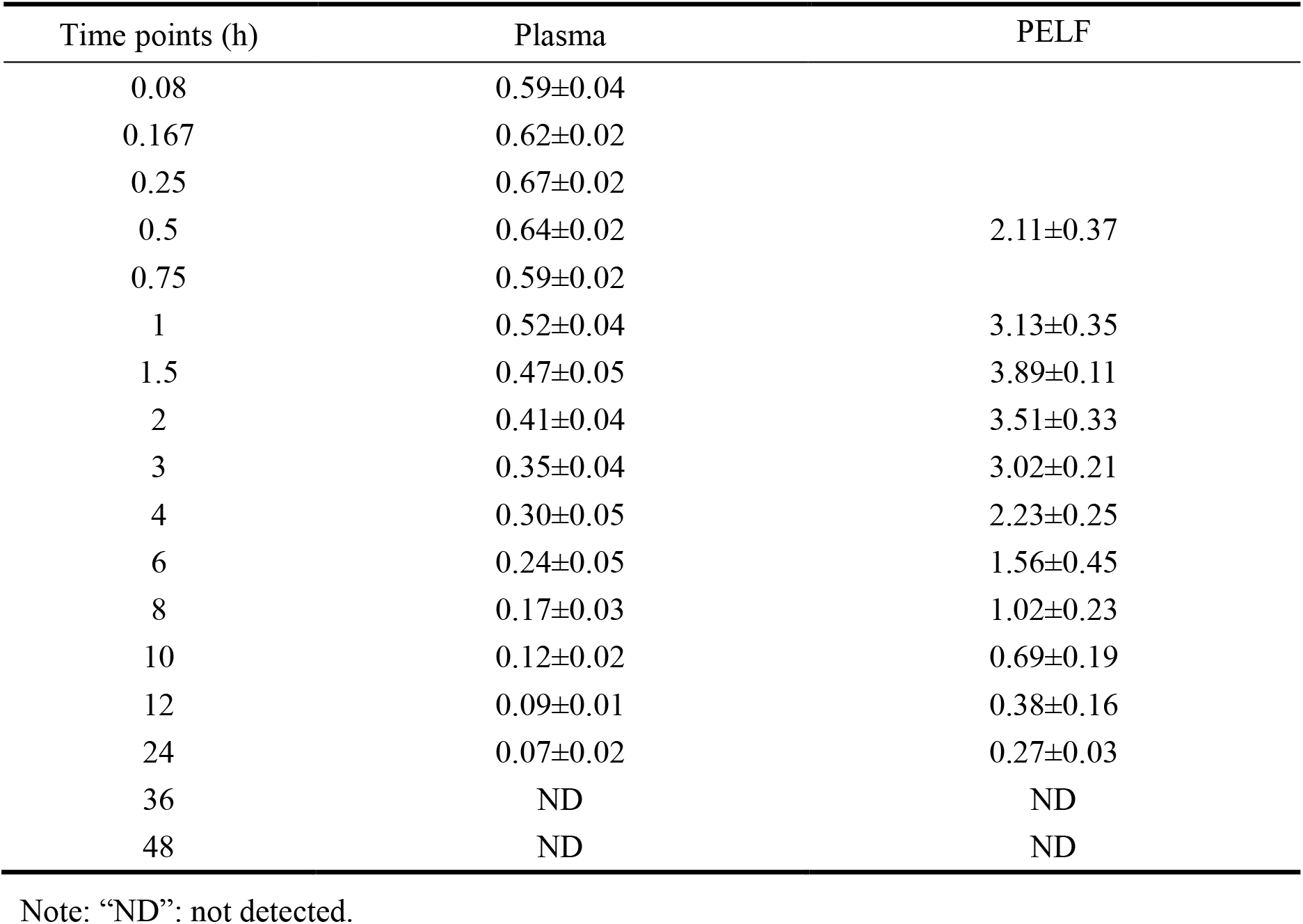
Concentrations of danfloxacin in plasma and PELF at various time points (n=6)

| Time points (h) | Plasma | PELF |
| --- | --- | --- |
| 0.08 | 0.59±0.04 |  |
| 0.167 | 0.62±0.02 |  |
| 0.25 | 0.67±0.02 |  |
| 0.5 | 0.64±0.02 | 2.11±0.37 |
| 0.75 | 0.59±0.02 |  |
| 1 | 0.52±0.04 | 3.13±0.35 |
| 1.5 | 0.47±0.05 | 3.89±0.11 |
| 2 | 0.41±0.04 | 3.51±0.33 |
| 3 | 0.35±0.04 | 3.02±0.21 |
| 4 | 0.30±0.05 | 2.23±0.25 |
| 6 | 0.24±0.05 | 1.56±0.45 |
| 8 | 0.17±0.03 | 1.02±0.23 |
| 10 | 0.12±0.02 | 0.69±0.19 |
| 12 | 0.09±0.01 | 0.38±0.16 |
| 24 | 0.07±0.02 | 0.27±0.03 |
| 36 | ND | ND |
| 48 | ND | ND |

Note: “ND”: not detected.

